# Discovering *in vivo* cytokine eQTL interactions from a lupus clinical trial

**DOI:** 10.1101/118703

**Authors:** Emma E. Davenport, Tiffany Amariuta, Maria Gutierrez-Arcelus, Kamil Slowikowski, Harm-Jan Westra, Yang Luo, Ciyue Shen, Deepak A. Rao, Ying Zhang, Stephen Pearson, David von Schack, Jean S. Beebe, Nan Bing, Sally John, Michael S. Vincent, Baohong Zhang, Soumya Raychaudhuri

## Abstract

**Background:** Cytokines are critical to human disease and are attractive therapeutic targets given their widespread influence on gene regulation and transcription. Defining the downstream regulatory mechanisms influenced by cytokines is central to defining drug and disease mechanisms. One promising strategy is to use interactions between expression quantitative trait loci (eQTLs) and cytokine levels to define target genes and mechanisms.

**Results:** In a clinical trial for anti-IL-6 in patients with systemic lupus erythematosus we measured interferon (IFN) status, anti-IL-6 drug exposure and genome-wide gene expression at three time points (379 samples from 157 individuals). First, we show that repeat transcriptomic measurements increases the number of *cis* eQTLs identified compared to using a single time point by 64%. Then, after identifying 4,818 cis-eQTLs, we observed a statistically significant enrichment of *in vivo* eQTL interactions with IFN status (p<0.001 by permutation) and anti-IL-6 drug exposure (p<0.001). We observed 210 and 72 interactions for IFN and anti-IL-6 respectively (FDR<20%). Anti-IL-6 interactions have not yet been described while 99 of the IFN interactions are novel. Finally, we found transcription factor binding motifs interrupted by eQTL interaction SNPs, pointing to key regulatory mediators of these environmental stimuli and therefore potential therapeutic targets for autoimmune diseases. In particular, genes with IFN interactions are enriched for ISRE binding site motifs, while those with anti-IL-6 interactions are enriched for IRF4 motifs.

**Conclusion:** This study highlights the potential to exploit clinical trial data to discover *in vivo* eQTL interactions with therapeutically relevant environmental variables.

## Background

Cytokines are critical signals used by the immune system to coordinate inflammatory responses. These factors bind to specific receptors to induce widespread transcriptional effects. Cytokines and their receptors are not only genetically associated with susceptibility to a range of human diseases, they have also emerged as effective therapeutic targets[1]. Blockade of tumour necrosis factor (TNF) was the first cytokine-directed therapy to achieve widespread use and is now used broadly to treat multiple inflammatory diseases including rheumatoid arthritis (RA), psoriasis, and inflammatory bowel disease[2]. More recently, IL-6 has emerged as a compelling therapeutic target. IL-6 levels are elevated in autoimmune diseases such as systemic lupus erythematosus (SLE) and RA. The IL-6 receptor has been successfully targeted with tocilizumab in RA[3] and giant cell arteritis[4], while IL-6 has been targeted directly with siltuximab for successful treatment of Castleman’s disease[5]. In SLE, IL-6 is thought to play a role in the observed B cell hyperactivity and autoantibody production[6]. Targeting IL-6-R in SLE has shown promise in phase I trials[7] and this has led to the development of other biologics targeting IL-6 such as PF-04236921[8]. Interferon (IFN)-α, produced primarily by plasmacytoid dendritic cells, has pleiotropic effects on the immune system. It has been implicated as a key mechanism in SLE development and pathogenesis, and is being investigated as a therapeutic target[9]. Agents targeting other inflammatory cytokines, including Interleukin-1 (IL-1), IL-12, and IL-17A and IL-23 are also in clinical use to treat autoimmune conditions. Interestingly, IL-1 blockade with canakinumab has also been recently reported to reduce risk of heart attacks, stroke and cardiovascular disease[10]. Therefore, defining the regulatory consequences of physiologic perturbations of cytokine levels will inform our understanding of both disease and drug mechanisms.

A *cis* expression quantitative trait locus (eQTL) contains a genetic variant that alters expression of a nearby gene. *Cis* eQTLs are ubiquitous across the genome[11] and while most are stable across tissues and conditions, environmental variables can alter the effects of some of them[12–18]. If an environmental change leads to disruption of regulators upstream of a gene, then it could magnify or dampen an eQTL effect, resulting in a genotype-by-environment interaction (**Figure S1**). Therefore, observing a set of eQTL interactions due to a perturbagen, such as a cytokine, can identify shared upstream regulatory mechanisms, such as transcription factors and key pathways. Even a single eQTL interaction where we can define mechanism can lead to insights about the action of the perturbagen.

However, *cis* eQTL interactions with physiologic environmental factors in humans have been challenging to discover *in vivo*[19–23] even with large cohorts[11, 17]. Success at finding *cis* eQTL interactions has largely been found in studies using model organisms[24, 25] or treating cells *in vitro* with non-physiologic conditions[26]. Thus far, these studies might be limited in power since they often map eQTLs separately across conditions and fail to exploit the power of repeat measurements[27]. In other instances, they test for genetic variants associated with differential expression and miss information about the magnitude of the eQTL effect in a specific condition[28].

We predicted that if the transcriptome is assayed at multiple time points under different exposure states, then the repeat measurements could lead to an increase in power to detect eQTLs and their interactions with environmental perturbations. If the same individual is assessed at multiple times, then the noise in transcriptomic measurements is reduced. Furthermore, repeat measurements from the same individuals when they are both unexposed and exposed to an environmental perturbagen allow for more accurate modelling of the effect of the perturbagen within those subjects.

Clinical trials, with their structured study design, may be the ideal setting to detect eQTL interactions with therapeutically important variables. In clinical trials, it is becoming increasingly common to collect transcriptional and genetic data alongside clinical and physiological data[29]. This extensive phenotyping of therapeutically important variables and biomarkers within the same individual at multiple time points provides a unique opportunity to identify *in vivo* eQTL interactions.

Here, we examined the modulation of eQTL effects by environmental factors that alter cytokine levels using data from a phase II clinical trial to evaluate the safety and efficacy of a neutralizing IL-6 monoclonal antibody (PF-04236921) in 157 SLE patients[8] (**Methods**). Many patients with SLE exhibit high levels of genes induced by type I IFN; these genes, known as the IFN signature, are a marker of disease severity[30, 31] and a pathogenic feature of SLE. This feature of the disease, together with exposure to anti-IL-6 leads to cytokine fluctuations in this cohort yielding opportunities to assess the impact of cytokine levels on eQTL effects. While this drug was not significantly different from placebo for the primary efficacy endpoint (proportion of patients achieving the SLE Responder Index (SRI-4) at week 24), biologically it effectively reduced free IL-6 protein levels (**Figure S2**). Given the key role of IL-6 and IFN in a range of diseases, the downstream regulatory effects of these cytokines are of great interest to study.

In this study, we leverage the power of repeat transcriptional and environmental measurements from a lupus clinical trial to identify *in vivo* eQTL interactions with IFN status and anti-IL-6 exposure. In the process, we define novel eQTL interactions for both IFN and IL-6.

## Results

We conducted whole blood high-depth RNA-seq profiling at 0, 12, and 24 weeks in anti-IL-6 exposed and unexposed individuals with the Illumina TruSeq protocol. We quantified 20,253 gene features and examined 1,595,793 genotyped and imputed common variants genome-wide (**Methods**). Along with each RNA-seq assay, we documented anti-IL-6 exposure and quantified IFN signature status with real-time PCR.

### Mapping eQTL in SLE patients

We first mapped *cis* eQTLs and then tested them for interactions with IFN status and anti-IL-6 exposure. eQTL interactions can be explored using our interactive visualisation tool (http://baohongz.github.io/Lupus_eQTL, **Figure S3**).

To identify *cis* eQTLs, we examined the association between gene expression and SNPs within 250kb upstream of the transcription start site and 250kb downstream of the transcription end site. In order to account for repeat measurements, with up to three RNA-seq assays per patient (Figure 1A, 379 samples from 157 patients, **Methods**), we used a linear mixed model. We included 25 gene expression principal components to maximise the number of eQTL detected and 5 genotyping principal components to account for the heterogeneity in ethnicity in our cohort (**Methods**). We observed that the multi-ethnic nature of our study did not confound our results, consistent with Stranger et al.[32] (**Figure S4**).

**Figure 1.**
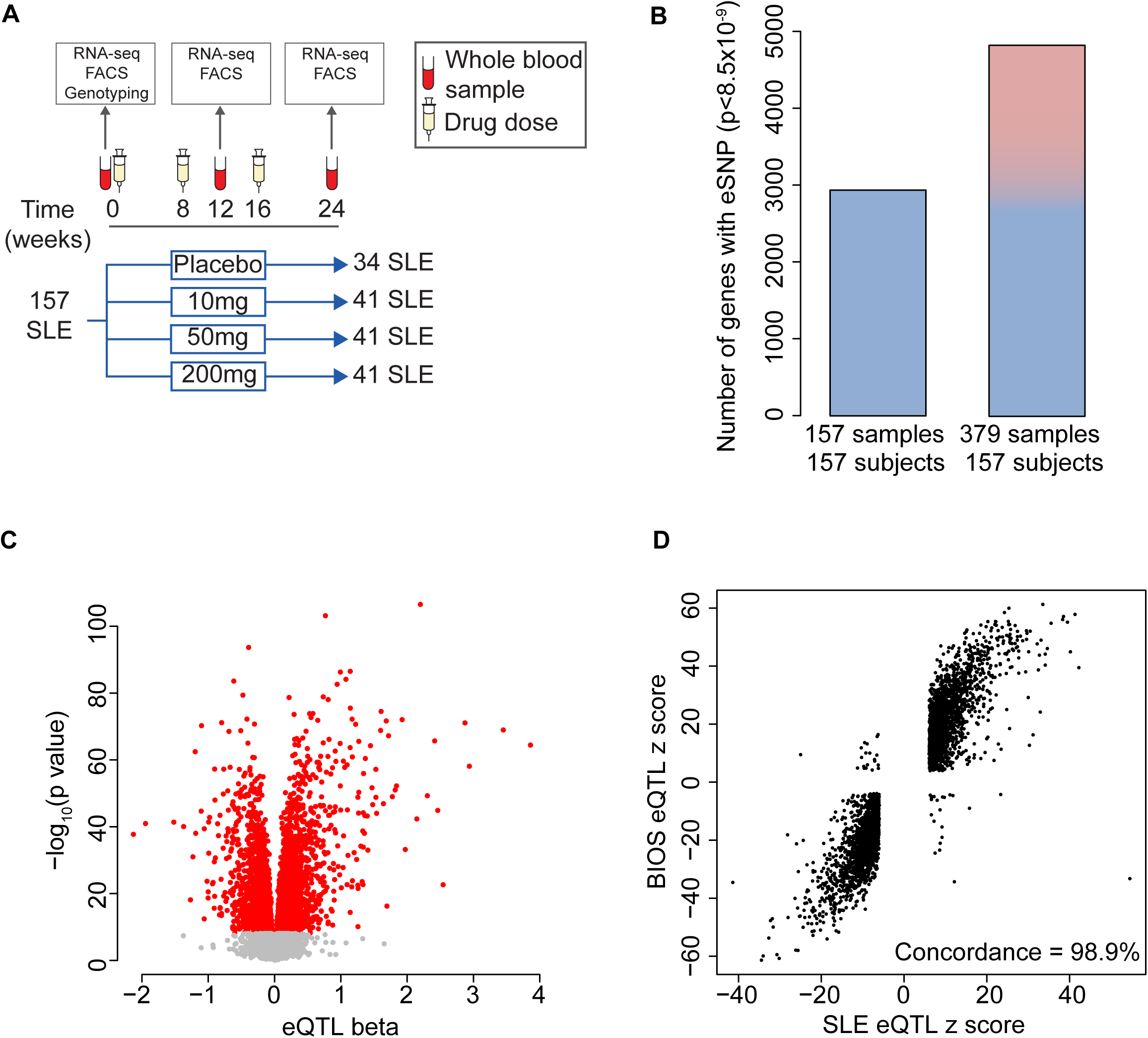
Identifying eQTLs in SLE patients. (A) Clinical trial structure and sampling strategy. (B) Number of eQTL genes identified using a linear model (left) and a linear mixed model (right). For the linear model, we used the first available time point for each individual (week 0 sample for n=152, week 12 sample for n=5). (C) Volcano plot of eQTL effects for the most significantly associated SNP for each gene (red color indicates p<8.5×10^−9^). (D) Concordance of SLE eQTL effects (p<8.5×10^−9^) with eQTLs observed in the BIOS cohort^11^ of healthy individuals (FDR <0.05). Each point represents the most significant SNP-gene pair for the SLE eQTL.

To ensure we only tested for interactions in a set of highly confident eQTLs, we applied a stringent correction for the total number of hypotheses tested. We recognized that this approach might arguably be overly stringent for eQTL discovery, but we wanted to be certain that we were only testing eQTLs for interactions that had a convincing main effect. Since we tested a total of 5,872,001 SNP-gene pairs genomewide, we set a significance threshold of p_eqtl_<8.5×10^−9^ (0.05/5,872,001 tests). We identified 4,818 *cis* eQTL genes (Figure 1B,1C, **Table S1**). The summary statistics for all the gene SNP pairs tested are available in **Table S2.**

To confirm the validity of our eQTLs, we compared them to a larger dataset. In the BIOS cohort, consisting of 2,166 healthy individuals[11], we observed that 85.4% of our SLE eQTL SNP-gene pairs are reported as eQTLs (FDR<0.05). Of these, 98.9% showed consistent direction of effect (p<5×10^−16^, binomial test, Figure 1D), suggesting that our results were highly concordant with those in this substantially larger study.

### Repeat measurements increase power to detect eQTL

Under reasonable assumptions, we would expect repeat samples to increase our power. Supporting that expectation, we detected 64% more *cis* eQTLs compared to the 2,934 genes from using a single sample (first available time point) per individual (Figure 1B). An alternative might have been to identify eQTLs separately from each of the three time points; however, this approach identified only a total of 3,050 eQTL genes (**Figure S5**). Modelling all three time points together results in 58% more *cis* eQTLs than modelling each time point separately.

We speculated that while repeat measures did increase power over single measures, that given a fixed number of samples, independent samples would lead to more power. To this end, we conducted an analysis fixing the number of samples at 157 and using 53 individuals with repeat measures (with two missing samples). Unsurprisingly, we found fewer eQTLs (2,215 genes) with the repeat measures alone compared to an analysis with the same number of independent samples (2,934 genes).

### IFN status eQTL interactions

For each of the 4,818 *cis* eQTL genes, we tested the most significantly associated SNP for environmental interactions with our linear mixed model framework. We first explored the influence of type I IFN on gene regulation after determining the IFN status of every patient at each time point. We classified each sample as either IFN high or IFN low using real-time PCR of 11 IFN-inducible genes[33] (**Methods**, Figure 2A).

**Figure 2.**
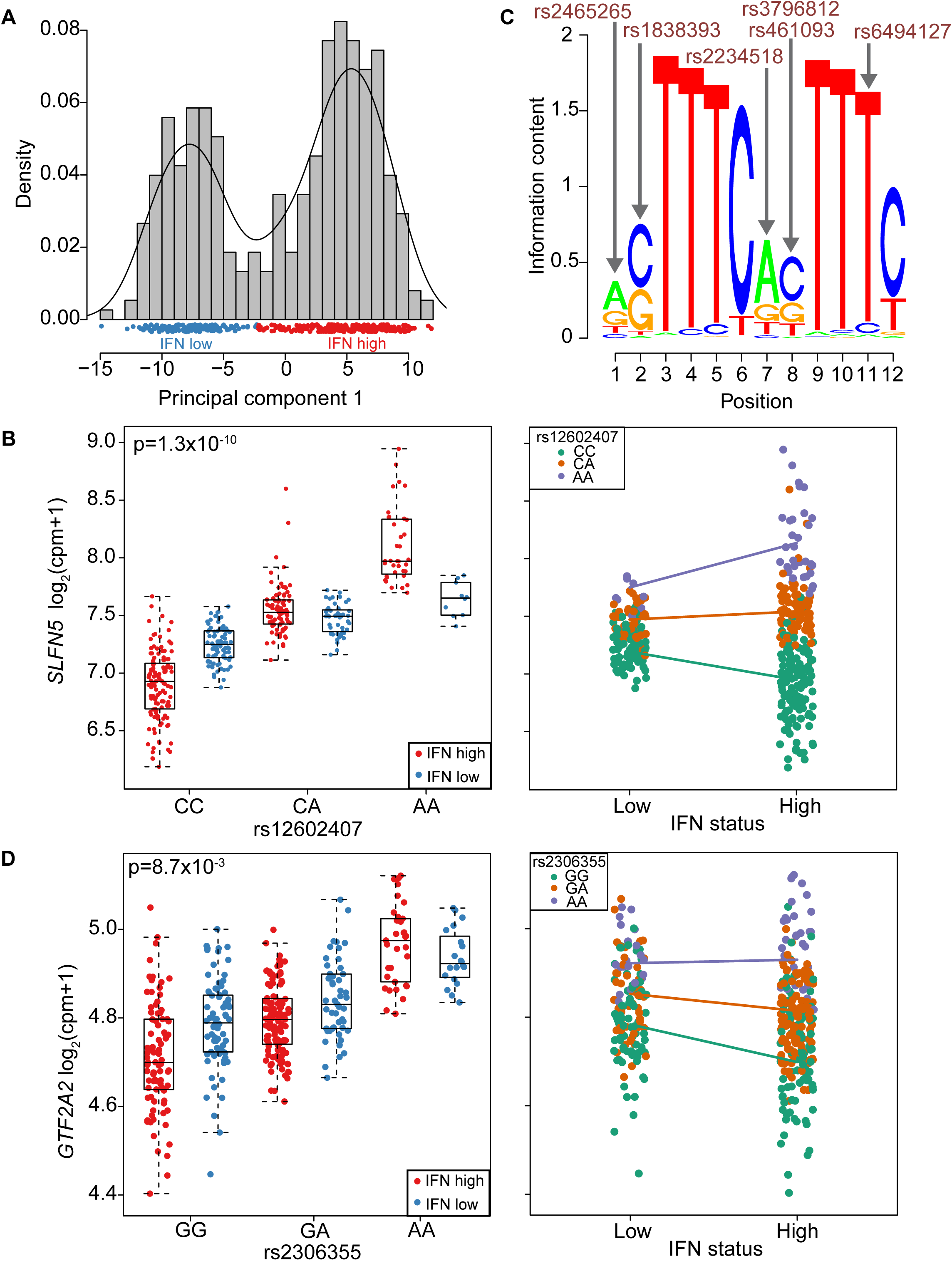
eQTL interactions with IFN status. **(A**) Designation of IFN status for each sample from the real-time PCR expression of 11 genes (first principal component). (**B**) IFN status interaction with the *SLFN5* eQTL plotted with respect to rs12602407 genotype (left) and IFN status of the sample (right). (**C**) The ISRE motif enriched among eQTLs magnified in IFN high samples. Arrows indicate positions of the motif interrupted by interaction SNPs (or SNPs in strong LD). Red indicates these SNPs correspond to magnified eQTLs. (**D**) IFN status interaction with the *GTF2A2* eQTL plotted with respect to rs2306355 genotype (left) and IFN status of the sample (right).

We first wanted to assess whether our results were indeed enriched for interactions. To do this, we identified those eQTLs with nominally significant interaction effects at p_interact_<0.01. We would expect ~48 out of 4,818 from chance alone. Surprisingly we observed 182 IFN-eQTL interactions (**Table S1**) that were nominally significant at p_interact_<0.01 suggesting that there was evidence of enrichment for eQTL interactions. We conducted permutations to ensure that these results were not the consequence of potentially inflated statistics, which might be the result for example of low frequency alleles, genes violating normality assumptions, or other technical artefacts. In each of 1,000 stringent permutations, we simply reassigned IFN status across samples and retested for eQTL interactions. This permutation preserves the main eQTL effect, since it maintains genotypes of the individuals with the associated expression data, but disrupts any real interactions that might be present in the data. In 0 out of 1,000 instance did we observe 182 or more interactions at p_interact_<0.01 suggesting that the number of observed interactions is enriched and highly unlikely to have happened by chance (**Figure S6**, p_permute_~0/1,000 = <0.001).

We then went on to identify those specific IFN-eQTL interactions of greatest interest by calculating a false discovery rate or q value for each interaction using the q value package[34] (**Methods**). We observed a total of 210 interactions with an FDR<0.2 threshold (**Table S1**). We note that 11 of these genes have already been described as having an interaction with a proxy gene for type I IFN signalling in the much larger BIOS study[11]. For example, *SLFN5* expression is influenced by the rs12602407 SNP (p_interact_ =1.3×10^−10^, FDR <9.9×10^−8^, Figure 2B) and this effect is magnified in IFN high samples. Of these 210 IFN-eQTL interactions, 99 were not reported in the BIOS study[11]. Indeed, applying a more stringent cut off of FDR<0.01, 27/34 of our interactions are not previously reported and therefore are almost certainly novel IFN-eQTL interactions with high confidence (**Figure S7**).

We speculated that groups of eQTL interactions might be driven by the same common regulatory factor. We divided interactions into magnifiers, where the environmental exposure increases the size of the eQTL effect, and dampeners where the environmental exposure decreases the eQTL effect (**Figure S8**). We hypothesized that the transcription factors driving the response to type I IFN may be different for the eQTL interactions defined as magnifiers (n=127, FDR<0.2) and dampeners (n=83, FDR<0.2).

We applied HOMER[35] to assess overlap between transcription factor binding motifs and the eQTL interaction SNPs (and SNPs in high linkage disequilibrium (LD, r^2^>0.8) in the *cis* window, Methods). We conducted two separate analyses: magnifying eQTL interactions in the foreground with dampening interactions in the background and vice versa. We found enrichment of motifs for key transcription factors involved in IFN signalling including a statistically significant enrichment for the ISRE motif (HOMER p=1×10^−4^, **Table S3**). The ISRE motif disruption occurred for 11 genes with an eQTL magnified in IFN high samples but for only 1 gene with an eQTL dampened (permutation p<0.019, **Methods**, Figure 2C). An example is the *GTF2A2* rs2306355 eQTL (p_interact_=8.7×10^−3^, FDR<0.15, Figure 2D); rs2306355 is in tight LD (r^2^=0.83 in Europeans) with rs6494127, which interrupts the TTCNNTTT core of the ISRE motif (Figure 3C). This SNP likely disrupts IRF9 and STAT2 binding in the ISGF3 complex[36], which binds to the ISRE motif. We observe greater expression of *GTF2A2* in individuals with the rs2306355 A allele compared to G; this difference is magnified in IFN high individuals (Figure 2D).

**Figure 3.**
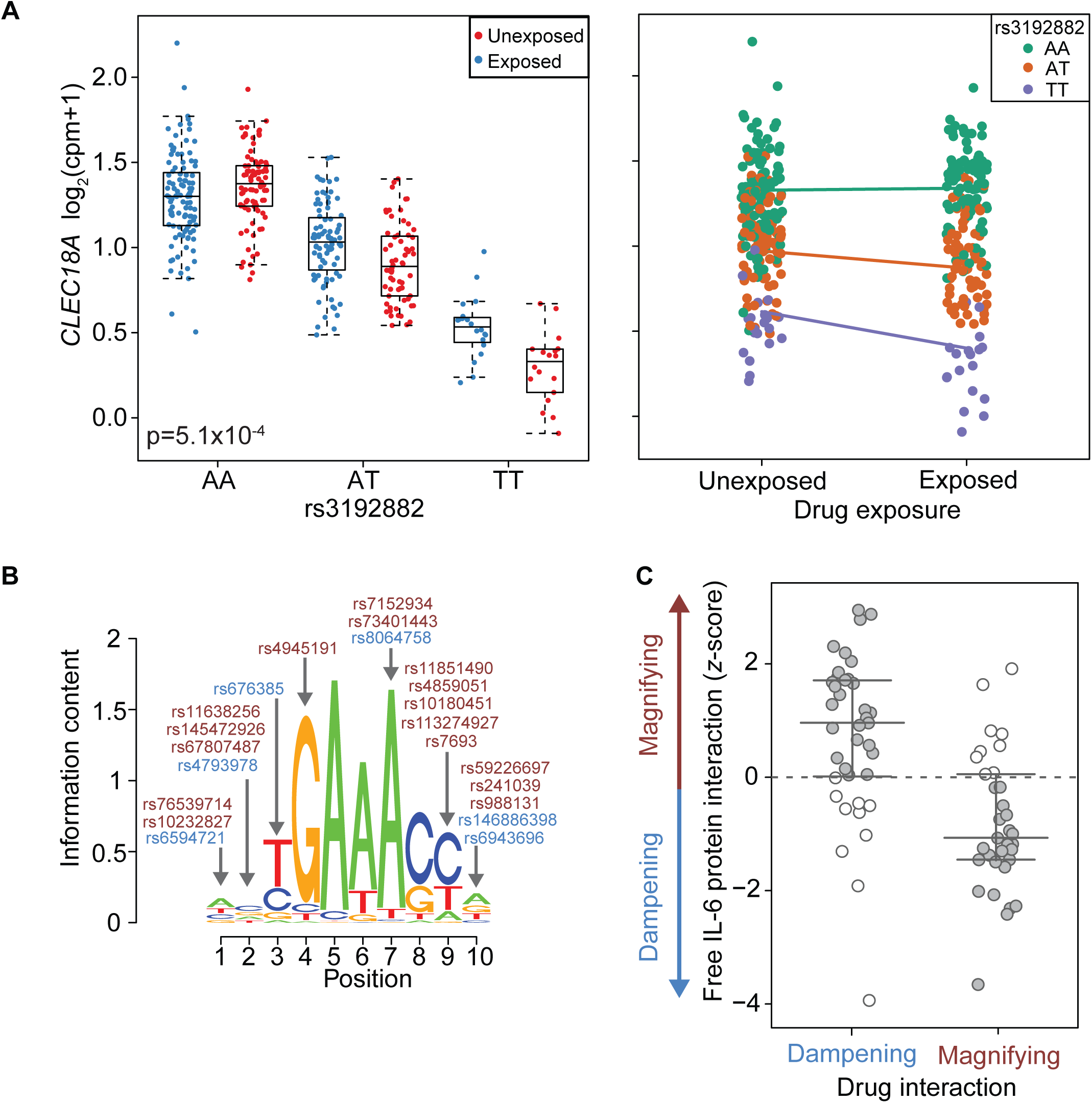
eQTL interactions with drug exposure. (A) Drug exposure interaction with the *CLEC18A* eQTL plotted with respect to rs3192882 genotype (left) and drug exposure (right). (B) The IRF4 motif enriched among eQTLs magnified following drug treatment. Arrows indicate positions of the motif interrupted by interaction SNPs (or SNPs in strong LD). Red and blue indicate SNPs corresponding to magnified and dampened eQTLs respectively. (C) Concordance of free IL-6 protein interaction effects with drug exposure interaction effects (grey indicates expected opposite direction).

We considered that the principal components included as covariates in our model might be mitigating power. For example, the 4^th^ principal component of gene expression is correlated with the IFN signature status of the sample (r_s_= −0.7, **Table S4**) so we repeated the IFN interaction analysis without correcting for principal component 4. For all the eQTLs tested for an IFN interaction, we observed very similar results with highly correlated z-scores (r_s_=0.94, **Figure S9**).

### Discovery of eQTL interactions with anti-IL-6 drug exposure

We then examined whether IL-6 blockade alters the relationship between genomic variation and gene expression and induces drug-eQTL interactions. We wanted to first test if there was evidence of such interactions in our data set. Again, using a threshold of p_interact_<0.01 for nominal significance for interactions, we observed 121 drug-eQTL interactions with anti-IL-6 out of 4,818 eQTLs tested (**Table S1**); similar to IFN interactions, this is far in excess of the ~48 we would expect by chance. As above, to ensure that these results were not the consequence of statistical artefact, we applied the same stringent permutation strategy, reassigning which samples were exposed or not to anti-IL-6. After 1,000 permutations, we never observed as many as 121 drug-eQTL interactions with p_interact_<0.01 (**Figure S10**), suggesting that our eQTLs were indeed highly enriched for those interacting with anti-IL-6 (p_permute_~0/1000<0.001).

To identify specific eQTL events that interact with anti-IL-6, we again calculated a false discovery rate. We observed that 72 of these interactions have an FDR<0.2. Only 8 of these drug-eQTL interactions overlap with the interactions observed for IFN status (**Table S1**). We note biologically relevant drug-eQTL interactions for *IL10* (p_interact_=2.6×10^−3^, FDR<0.19, **Figure S11**), an anti-inflammatory cytokine, *CLEC4C* (p_interact_=2.9×10^−3^, FDR<0.19) which has previously been associated in *trans* with an SLE risk allele[37] and *CLEC18A* (p_interact_=5.1×10^−4^, FDR<0.14, Figure 3A) another member of the C-type lectin domain family.

Similar to the IFN-eQTL interactions, we divided the drug-eQTL interactions into magnifiers (n=33, FDR<0.2) and dampeners (n=39, FDR<0.2) (**Figure S12**) and used the approach as described above to define transcription factors potentially driving the response to IL-6 blockade (**Table S5**). One of the motifs enriched for eQTLs magnified after drug treatment (with dampeners in the background) was IRF4 (HOMER p=1×10^−3^). The IRF4 motif disruption occurred for 9 genes, including *CLEC18A*, with an eQTL magnified after drug treatment compared to 4 genes with an eQTL dampened (Figure 3B, **Methods**). We permuted the magnifying and dampening genes and found this ratio for enrichment is suggestive at the gene level (p=0.058) but additional eQTL interactions will be necessary to confirm.

### Comparing differential expression to eQTL interactions

A more common strategy to determine the effect of an environmental variable is to use differential gene expression. For differential expression following anti-IL-6 treatment, we identified 415 genes with FDR<0.05 but modest effects (max fold change=1.3, **Figure S13**). Intriguingly, only 1/72 drug-eQTL interaction genes also show evidence of differential gene expression. This suggests that eQTL interactions offer independent information from differential expression, which might contribute to defining mechanisms.

### Concordance of drug-eQTL interactions with protein level interactions

We hypothesized that interactions due to drug exposure are likely driven by free IL-6 cytokine levels (our key clinical biomarker of interest). If this is the case, for eQTLs dampened by drug exposure, an increase in free IL-6 should elicit an opposite interaction effect and result in eQTL magnification. We assessed whether eQTL interactions with free IL-6 protein levels measured in the patient serum samples were consistent with those following IL-6 blockade. We observed enrichment in the overlap between cytokine interactions and drug interactions (53/72 interactions in expected opposite direction, Figure 3C, p=3.8×10^−5^, binomial test).

## Discussion

In this study we mapped eQTLs in a clinical trial of SLE patients and discovered interactions with IFN and IL-6, two clinically important cytokines. Our study had dramatic variation in IL-6 that was therapeutically induced, and variation in IFN due to the disease status of the SLE patients. This, together with the structured study design with repeat measurements of gene expression across different conditions in the same individual, allowed us to identify *in vivo* eQTL interactions.

eQTL interactions with drug interventions or other therapeutically relevant physiologic variables are important to identify. We note that these results are independent of differentially expressed genes, which are more likely to represent second order effects, rather than primary genetic effects. Therefore, these eQTL interactions can point to regulatory mechanisms, such as transcription factors or subclasses of enhancers, acting downstream of the environmental condition of interest and driving groups of eQTL interactions. The IFN status eQTL interactions we identified provide support for this approach. By making use of the direction of effect for the eQTL interaction, we were able to identify an enrichment of magnifying eQTL interaction SNPs interrupting the binding sites of transcription factors known to be important in the response to IFN, such as ISGF3 (the STAT1, STAT2 and IRF9 complex), which binds ISRE. Once we are able to recognize the downstream drivers of therapeutically relevant clinical variables, then it may become possible to define more mechanisms of action for drugs and more precise drug targets.

As a powerful example, we note enrichment of magnifying anti-IL-6-eQTL interaction SNPs interrupting the binding site of IRF4. It has been suggested that IRF4 works downstream of IL-6 by binding BATF and co-ordinately regulating the production of IL10 and other genes[38]. Consistent with this, we observed that the *IL10* eQTL does indeed interact with presence of anti-IL-6 (**Figure S11**). Previous studies have highlighted a role for IRF4 in the pathogenesis of autoimmune diseases in mouse and humans. For example in a murine model of SLE, *IRF4* knockout mice did not develop lupus nephritis[39]. In humans, IRF4 is associated with RA[40], a disease in which anti-IL-6 treatment has been successful[3]. Our findings provide further support that IRF4 could be a potential therapeutic target for autoimmune diseases such as RA where anti-IL-6 is effective[41].

The ability to focus on interactions with specific patient phenotypes might point to key targets for disease intervention. For example, IFN is a key immunophenotype in SLE patients, and elevated in SLE compared to healthy controls[30, 31]. The IFN status immunophenotype is already itself driving interest in therapeutic targets. A recent phase II clinical trial has shown that an antagonist to the type I IFN receptor, acting upstream of ISRE, reduced severity of symptoms in SLE. Interestingly, the antagonist was more effective in the patients with a high baseline IFN status[42]. This example provides a compelling case study for how understanding master regulators of key disease phenotypes might lead to promising new therapeutic strategies. We speculate that this provides a mechanism for stratified medicine for future studies, which may be applicable to other diseases.

We recognized that computing eQTL interactions requires a robust statistical model that accounts for genotype, environmental factor, RNA expression levels, repeat measurements, and technical covariates. We were sensitive to the possibility that pre-processing and normalisation of these factors could potentially have an impact on our results. For this reason, we used stringent filtering and examined only variants that were common and where the minor allele was present for each of the exposure groups. Next, to confirm enrichment of eQTL interactions, we used a stringent permutation-based strategy that preserved the distribution of genotypes and corresponding expression values. Finally, we also utilized a standard normal transformation[43] (**Methods**) and observed that this had little effect on the primary eQTL analysis (r_s_=0.99 for z scores, **Figure S14**) and interaction analyses (IFN r_s_=0.84, drug r_s_=0.76 for z scores, **Figure S15**), or the observed enrichment over the null in our stringent permutation analysis (**Figure S16**).

We speculate that drug-eQTL interactions might offer an alternative pharmacogenetic strategy to assess drug response. For many biologic medications, predictive pharmacogenetics through typical association studies has been challenging; for example, studies trying to define genetic or transcriptomic biomarkers of anti-TNF response have not been successful[44, 45]. An eQTL interaction approach can be used to define a genotype-aware score reflecting the biological activity that a medication is having upon an individual, given their allelic combination of multiple genetic markers. For example, we can define a simple anti-IL-6 exposure score based on 7 anti-IL-6 eQTL interactions with a more stringent FDR (FDR<0.1). This score is based on assessing whether the expression of the eQTL target gene was more consistent with the drug exposed or the unexposed state for the corresponding interaction SNP genotype. Unsurprisingly, we found a difference in drug exposure score between the unexposed and exposed samples (**Figure S17**) (r_s_=0.40, p=2.1×10^−16^); these differences reflect the fact that the eQTLs were themselves identified by examining samples with and without drug exposure. However, while we did not utilize the administered drug dose to identify drug-eQTL interactions; we observed a significant correlation between drug dose (10, 50 or 200mg) and drug exposure score (r_s_=0.16, p=0.02) in the drug-exposed samples (**Figure S18**). A simple eQTL interaction score may therefore have the potential to stratify individuals when assessing response to a medication for example those with a higher drug exposure score may have a better response to treatment. Similarly, this score could be correlated with adverse effects to capture informative gene expression signatures.

We note a limitation of this study is that the drug itself did not achieve its primary efficacy endpoint of improving SLE outcomes. Hence, while the drug exposure score for this study tracked with the biological effect of the drug (reducing free IL-6 protein levels), it might not be useful for SLE specifically. However, such a scoring system could be implemented easily in most phase III trials for a broad range of therapeutics, where the numbers of samples are far in excess of this phase II trial, ensuring better powered and more accurate eQTL-interaction mapping.

## Conclusion

We devised a framework for identifying *in vivo* eQTL interactions with therapeutically relevant variables, exploiting repeat measurements from a clinical trial. We have applied this approach to demonstrate how downstream regulatory effects of cytokine biology can be elucidated. This same approach can be applied to a wide range of other clinically important cytokines, their antagonists, or indeed other targeted biologic therapies. We speculate that this approach might even be applied to the presence or absence of disease, or disease activity. However, given the multifaceted nature of disease effects, interpreting an eQTL interaction in that context might be more challenging. Modern clinical cohorts and clinical trial data sets with RNA-seq data that has been collected will make this approach easily applicable on a wide scale.

## Methods

### Study design

The objectives of this study were to map eQTLs in a cohort of lupus patients and identify eQTL interactions with environmental perturbations such as drug treatment to shed light on drug and disease mechanisms. SLE patients were recruited to a phase II clinical trial to test the efficacy and safety of an IL-6 monoclonal antibody (PF-04236921). The patient population recruited to this trial have been detailed extensively by Wallace et al.[8] 183 patients (forming a multi-ethnic cohort) were randomized to receive three doses of drug (10, 50 or 200mg) or placebo at three time points during the trial (weeks 0, 8 and 16).

### RNA-sequencing

We collected peripheral venous blood samples in PAXgene Blood RNA tubes (PreAnalytiX GmbH, BD Biosciences) for high-depth RNA-seq profiling at 0, 12, and 24 weeks. We extracted total RNA from blood samples using the PAXgene Blood RNA kit (Qiagen) at a contract lab using a customized automation method. We assessed the yield and quality of the isolated RNA using Quant-iT^™^ RiboGreen^®^ RNA Assay Kit (ThermoFisher Scientific) and Agilent 2100 Bioanalyzer (Agilent Technologies), respectively. Following quality assessment, we processed an aliquot of 500-1000 ng of each RNA with a GlobinClear-Human kit, (ThermoFisher Scientific) to remove globin mRNA. We then converted RNA samples to cDNA libraries using TruSeq RNA Sample Prep Kit v2 (Illumina) and sequenced using Illumina HiSeq 2000 sequencers. We generated an average of 40M 100bp pair-end reads per sample for downstream analysis.

We successfully obtained 468 RNA-seq profiles from 180 patients. We aligned reads to the reference genome and quantified gene expression using Subread[46] and featureCounts[47] respectively. We included genes with at least 10 reads (CPM>0.38) in at least 32 samples (minimum number of patients with both unexposed and exposed RNA-seq assays in a drug group) prior to normalization. Following quality control (QC), we removed 4 samples as outliers. We then normalized 20,253 transcripts using the trimmed mean of M-values method and the edgeR R package[48]. Expression levels are presented as log_2_(cpm +1) (**Table S6**).

### Genotyping

We genotyped 160 individuals across 964,193 variants genome-wide with the Illumina HumanOmniExpressExome-8v1.2 beadchip. We removed SNPs if they deviated from Hardy-Weinberg Equilibrium (HWE) (p < 1×10^−7^), had a minor allele frequency <5%, missingness >2% or a heterozygosity rate greater than 3 standard deviations from the mean (PLINK[49, 50]). For mapping eQTLs, we removed SNPs on the Y chromosome. Following QC, we used 608,017 variants for further analysis. We removed one sample with high missingness and outlying heterozygosity rate from further analysis.

### Imputation

We pre-phased the genotypes with SHAPEIT v2[51]. We imputed missing genotypes and untyped SNPs using Impute2[52] in 5Mb chunks against the 1000 Genomes Phase 3[53] reference panel. To ensure only highly quality genotypes, and to avoid artefacts that can be induced by imputation uncertainty, we removed SNPs with an info score <1, MAF<0.05 or HWE p<1×10^−7^ leaving 1,595,793 SNPs for further analysis.

### Interferon status

We classified the interferon (IFN) status of each sample at each time point from the expression of 11 IFN response genes (*HERC5*, *IFI27, IRF7, ISG15, LY6E, MX1, OAS2, OAS3, RSAD2, USP18, GBP5)* using TaqMan Low Density Arrays. These 11 genes were selected by identifying transcripts for which there was both a measureable response to IFN treatment *in vitro*, as well as differential expression (reduction in expression level) between baseline and visits with clinical improvement in the BOLD study[33]. There is no consensus set of genes to determine the IFN status of SLE patients but these 11 genes do overlap with other published gene sets. For example 4/11 genes are also used in the 7-gene set defined by McBride et al[54] and 9/11 genes overlap with the 21-gene set defined by Yao et al[55].

The first principal component of the expression of the 11-gene set captured 91.7% of the variation (**Figure S19**). The distribution of this first principal component is nearly bimodal with good separation (Figure 2A) and we classified samples as high or low IFN based on this first principal component score. In our dataset, we see excellent correlations (r_s_=0.86-0.98) between the real-time PCR expression and the RNA-seq expression for these 11 genes (**Figure S20**). The first PC of the IFN signature of RNA-seq data is also strongly correlated with the first PC of the IFN signature of real-time PCR (r_s_=0.96, **Figure S21**). IFN status was available for 376 samples from 157 subjects.

### Drug exposure

Samples were assigned as unexposed (placebo or week 0 samples) or drug exposed (week 12 and week 24 samples in the drug groups).

### Free IL-6 protein levels

We determined free IL-6 protein levels from serum using a commercial sandwich ELISA selected for binding only free IL-6. The assay was validated according to FDA biomarker and fit-for purpose guidelines. Free IL-6 protein levels were available for 311 samples from 145 subjects. Since the distribution of IL-6 levels was highly skewed, we ranked samples in order of IL-6 protein levels and included in the model to identify drug-eQTL interactions.

### Statistical analysis

#### eQTL and interaction analysis

In total, 157 patients (with 379 RNA-seq samples) had good quality gene expression and genotyping data for eQTL analysis. All statistical analyses were carried out in R[56].

We defined a *cis* eQTL as the SNP within 250kb upstream of the GENCODE[57] transcription start site of the gene or 250kb downstream of the transcription end site. We first applied a linear model for the first available time point (week 0 sample for n=152, week 12 sample for n=5) to identify each eQTL using the first 25 principal components of gene expression and the first 5 principal components of genotyping as covariates.

To select the number of gene expression principal components to include, we counted the number of eQTL genes identified after incrementally increasing the number of principal components accounted for in the model from 0 to 50 by increments of five (**Figure S22**). We selected 25 principal components of gene expression to maximise the number of eQTL genes detected while minimising the number of principal components we corrected for. We included 5 principal components of genotyping to account for the heterogeneity in ethnicity in our cohort (**Figure S23**).

SNPs were encoded as 0, 1 and 2 with respect to the number of copies of the minor allele. To adjust for multiple testing during eQTL discovery we used a stringent Bonferroni corrected *p*-value threshold of 8.5×10^−9^ (0.05/ 5,872,001 tests). The Bonferroni adjustment assumes independence among the tests and we therefore note that it is a conservative multiple comparisons adjustment.

To map eQTLs using multiple samples for each individual, we applied a random intercept linear mixed model using the first 25 principal components of gene expression and the first 5 principal components of genotyping as covariates and patient as a random effect:

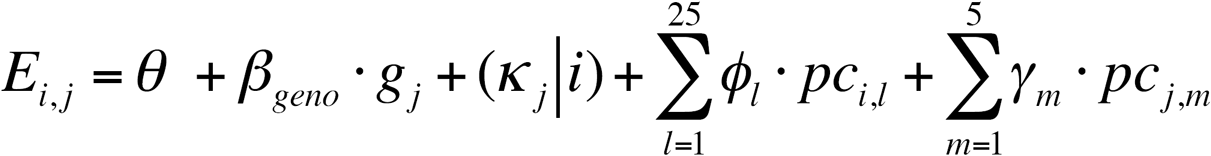

Where *E_i,j_* is gene expression for the *i^th^* sample from the *j^th^* subject, *θ* is the intercept, *β_geno_* is the genotype effect (eQTL), *(κ_j_|i)* is the random effect for the i^th^ sample from the j^th^ subject, *pc_i,l_* is principal component l of gene expression for sample i, *pc_j,m_* is principal component *m* of genotyping for subject *j*.

We fitted the linear mixed models using the lme4 R package[58]. We assumed covariance between samples from the same individual, but did not assume any structure in this covariance.

We used the most significant SNP (with p<8.5×10^−9^) from the 4,818 identified eQTL genes to explore eQTL interactions. For each environmental interaction analysis, we further filtered these eQTLs to include only those with at least two individuals homozygous for the minor allele of the SNP being tested in each of the environmental factor groups. For example, we required two of these individuals in each of the drug exposed and drug unexposed groups. To identify eQTL interactions, we added an additional covariate to the model for example drug exposure, and an interaction term between this covariate and the genotype of the SNP:

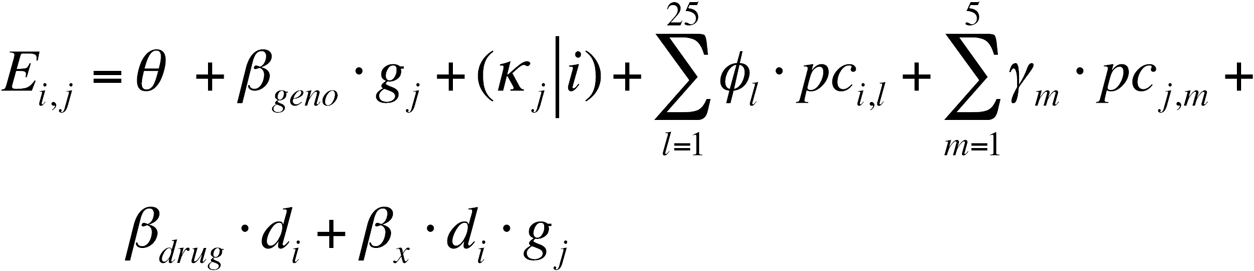

Where *E_i,j_* is gene expression for the *i^th^* sample from the *j^th^* subject, *θ* is the intercept, *β_geno_* is the genotype effect (eQTL), *(κ_j_|i)* is the random effect for the i^th^ sample from the j^th^ subject, *pc_i,l_* is principal component l of gene expression for sample i, *pc_j,m_* is principal component *m* of genotyping for subject *j, β_drug_* is the drug effect (differential gene expression) and *β_x_* is the interaction effect.

We determined the significance of the interaction term with a likelihood ratio test.

To rigorously confirm the relative enrichment of eQTL interactions, we shuffled the interaction covariate (for example drug exposure) 1,000 times and calculated the number of significant interactions observed in each permutation. Our primary goal for the permutation analysis was to retain the main eQTL effect while examining only the effect of the environmental factor on the interaction. In this study, the main purpose of the covariates included in the model is to ensure the main eQTL effect is found. For IFN high/low status, we shuffled across all samples. For drug interaction permutation analysis, we maintained the number of individuals in the drug group and the number of samples with exposure to drug. We calculated a qvalue for each interaction using the q value package[34]. **Figure S24** shows the observed versus the expected p values for the interaction analyses.

The expression of the majority of genes followed a normal distribution (**Figure S25**) but to assess whether non-normality could be causing an inflation of our test statistic, we repeated the identification of eQTLs and eQTL interactions following the standard normal transformation. We transformed the expression values of each gene to their respective quantiles of a normal distribution using the qqnorm function in R, breaking any ties (for example expression levels of zero in some individuals) randomly.

### Concordance with an eQTL study in healthy individuals

In the SLE cohort, we classified 4,818 *cis* eQTL genes (p<8.5×10^−9^). The z-score for the most associated SNP for each of these genes was compared to the z-score from a previously published eQTL dataset from whole blood from 2,166 healthy individuals[11]. 4,113/4818 SNP-gene pairs (85.4%) were also reported in the BIOS dataset (FDR<0.05). After removing 301 SNPs, which could not be mapped to a strand 3,770/3,812 (98.9%) had a z-score (eQTL effect) in a consistent direction.

### Magnifiers and Dampeners

An eQTL interaction can either magnify or dampen the original eQTL effect. We multiplied the interaction z-score by the sign of the original eQTL effect (genotype beta) and defined magnifiers as interactions with an adjusted z-score > 0 and dampeners as interactions with an adjusted z-score < 0.

### Differential gene expression analysis

To identify differentially expressed genes following drug exposure (unexposed or exposed), we applied a random intercept linear mixed model using the first 25 principal components of gene expression and the first 5 principal components of genotyping as covariates and patient as a random effect. We calculated a q value using the q value package[34].

### Drug exposure score

We assigned a drug exposure score to each sample. We calculated a score for each gene (see equation below) and then averaged across the 7 drug-eQTL genes (FDR<0.1) to give the final drug exposure score.

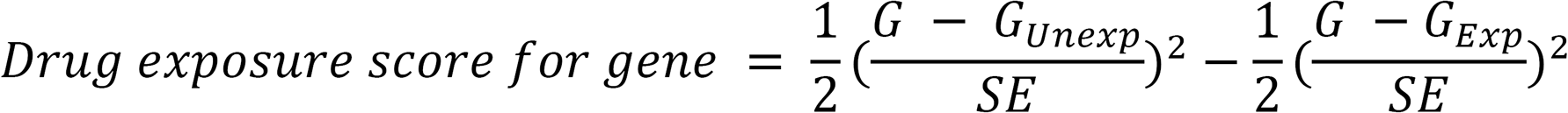

Where G is gene expression for a given sample, G_Unexp_ is predicted mean gene expression for unexposed samples of the relevant SNP genotype, G_Exp_ is predicted mean gene expression for exposed samples of the relevant SNP genotype and SE is standard error for the intercept term of the model (unexposed expression for genotype 0).

### HOMER analysis for transcription factor binding motif enrichment

We used the HOMER software suite[35] to look for enrichment of transcription factor binding motifs in the 210 IFN-eQTL interactions (FDR<0.2) and the 72 drug-eQTL interactions (FDR<0.2). Each eQTL interaction was identified using the most highly associated SNP for that eQTL. However, as this SNP is not necessarily the functional SNP, we additionally considered all those with an r^2^≥0.8 in the 1000 Genomes European population[53] within the *cis* eQTL window. We defined our motif search window as 20 bp on either side of each SNP (i.e. 41 bp wide).

For each environmental factor, we divided the eQTL interactions into magnifiers or dampeners and conducted two separate HOMER analyses: one with magnifiers in the foreground and dampeners in the background; the other with dampeners in the foreground and magnifiers in the background. HOMER reported the transcription factor motifs that were significantly enriched in the foreground relative to background. Motifs were plotted using the SeqLogo R library[59].

We determined permutation p values for enrichment of the ISRE and IRF4 transcription factor binding sites as follows. For ISRE, the motif is interrupted by interaction SNPs (or SNPs in LD) corresponding to 11 magnifying genes and 1 dampening genes. We permuted which genes were labelled as magnifiers or dampeners 100,000 times and counted the number of genes in each category with an ISRE motif interrupted. We found 1,855 occurrences from 100,000 trials with at least 11 magnifying genes (p<0.019). For IRF4 the motif is interrupted by SNPs corresponding to 9 magnifying genes and 4 dampening genes. Using the same permutation approach, we found 5,801 occurrences from 100,000 trials with at least 9 magnifying genes (p<0.058).

## Declarations

### Ethics approval and consent to participate

The protocol was approved by each institutional review board subject to applicable laws and regulations and ethical principles consistent with the Declaration of Helsinki.

### Availability of data and material

Data are available in the supplementary tables and at http://baohongz.github.io/Lupus_eQTL.

## Acknowledgements

This work is supported in part by funding from the National Institutes of Health (U01GM092691, UH2AR067677, U19AI111224 (SR)), the Doris Duke Charitable Foundation Grant #2013097, the Ruth L. Kirschstein National Research Service Award (F31AR070582) from the National Institute of Arthritis and Musculoskeletal and Skin Diseases (KS) and the Rheumatology Research Foundation Tobé and Stephen E. Malawista, MD Endowment in Academic Rheumatology (D.A.R). This work is also supported by unrestricted funding from Pfizer, Inc.

## Author Contributions

The project was conceived and designed by EED, MSV, BZ and SR. Statistical analysis was conducted by EED, TA, MG-A, KS, H-JW, YL and CS. Molecular data was obtained, organized and analysed by YZ, SP, DvS, JSB, NB, MSV, BZ and DAR. The initial manuscript was written by EED and SR. All authors edited and approved the manuscript.

